## Supplemental information for "Fundamental bounds on learning performance in neural circuits"

February 15, 2019

### Supplementary notes on mathematical analyses

#### Necessary Conditions on Learning

In this section, we specify some necessary requirements on any learning rule, and provide insight into factors that make a task 'easier' to learn. We consider a situation in which a neural network has to learn a task, by changing its synaptic strengths (represented in a vector $\mathbf{w}$). The level of training is represented by some task-error function $F[\mathbf{w}]$, which decreases with increasing levels of performance. In order to learn, the network receives some form of sensory feedback on task error, which informs changes at each synapse through some unknown learning rule.

We will first ask what it takes for a network to learn, given limited rates of sensory feedback on task error, over some time interval $[0, T]$. Clearly, learning is equivalent to a decrease in task error, which corresponds to

$$F[\mathbf{w}(T)] - F[\mathbf{w}(0)] < 0.$$

The degree of learning is determined by how much smaller than zero this quantity is. Suppose our task error is continuously differentiable in $\mathbf{w}$. Then, by the Fundamental Theorem of Calculus,

$$\begin{aligned} F[\mathbf{w}(T)] - F[\mathbf{w}(0)] &= \int_0^T \langle \nabla F[\mathbf{w}(t)], \dot{\mathbf{w}}(t) \rangle \; dt \\ &= T \; \mathbb{E}_t[\langle \nabla F[\mathbf{w}(t)], \dot{\mathbf{w}}(t) \rangle], \end{aligned} \tag{S.1}$$

where expectation is taken across a uniform distribution of timepoints on the interval $[0, T]$, and $\dot{\mathbf{w}}(t)$ denotes the time-derivative of $\mathbf{w}$. Let us pick (uniformly) a random synaptic weight $w_i$, at a (uniformly) random timepoint $t \in [0, T]$. If learning has occurred, then the above equations imply

$$\mathbb{E}\big[\nabla F[\mathbf{w}(t)]_i, \dot{w}_i(t)\big] < 0.$$

In other words, at time $t$, we expect the average synapse to be changing in a direction that anticorrelates with $\nabla F[\mathbf{w}(t)]_i$, the $i^{th}$ component of the gradient of task performance at time $t$. Therefore, any learning rule must possess (on average) information on $\nabla F[\mathbf{w}(t)]$ at time $t$. However, any learning rule will be using 'old' information at time $t$. Information on task error may only be supplied intermittently, and there must exist some biochemically induced delay between sensory acquisition of information on task error, and its integration into plasticity rules at each synapse. How well a learning rule performs therefore depends on how relevant 'old' information is.

We can obtain intuition on the previous assertion. Suppose the plasticity direction $\dot{\mathbf{w}}(t)$ was determined using 'old' information from time 0. Suppose that the learning rule calculated an estimate $\nabla \tilde{F}[\mathbf{w}(0)]$ of $\nabla F[\mathbf{w}(0)]$. If the gradient was constant over the time interval $[0,t]$, then we would have $\nabla F[\mathbf{w}(0)] = \nabla F[\mathbf{w}(t)]$, and the delay from time 0 to time $t$ would have no effect on the quality of the estimate. Conversely if the gradient changed very fast over this period, then the delay would greatly decrease the quality of the estimate. What influences how fast the gradient's direction changes? Geometrically, the rate of change of the gradient is the curvature, or second derivative, of $F[\mathbf{w}]$. Thus the geometry of $F[\mathbf{w}]$ is one factor. Another is the overall speed of synaptic change: if the network moves faster through weight space, then the gradient direction will also change faster. Task irrelevant plasticity from synaptic processes unconnected with learning also contributes to this speed, and therefore hinders learning in general.

One of the results in the paper is a demonstration of how increasing network size can 'flatten out' an error function $F[\mathbf{w}]$. This flattening makes any learning rule with unavoidable amounts of both delay between information acquisition and synaptic plasticity, and task-irrelevant plasticity, learn faster.

### Learning rate and local task difficulty

In order to quantify learning rate over the time interval $[0,T]$, we take $k$ such that

$$F[\mathbf{w}(T)] = [1 - kT]F[\mathbf{w}(0)].$$

Meanwhile we represent overall synaptic change over the time interval $[0,T]$, normalised by $T$, as

$$\dot{\boldsymbol{\omega}}_T = \frac{\mathbf{w}(T) - \mathbf{w}(0)}{T}.$$

We perform a second order Taylor expansion of the above equation to get:

$$-kTF[\mathbf{w}(0)] = F[\mathbf{w}(T)] - F[\mathbf{w}(0)] \tag{S.2a}$$

$$= T\langle \nabla F[\mathbf{w}(0)], \dot{\boldsymbol{\omega}}_T \rangle + \frac{1}{2}T^2 \langle \dot{\boldsymbol{\omega}}_T, \nabla^2 F[\mathbf{w}(0)] \dot{\boldsymbol{\omega}}_T \rangle + \mathcal{O}(T^3). \tag{S.2b}$$

We can rearrange this equation to make $k$ the subject. This gives

$$k = \frac{-\|\nabla F[\mathbf{w}(0)]\|_2}{F[\mathbf{w}(0)]} \left[ \left\langle \dot{\boldsymbol{\omega}}_T, \nabla \hat{F}[\mathbf{w}(0)] \right\rangle + \mathbf{G}_F[\dot{\boldsymbol{\omega}}_T] \|\dot{\boldsymbol{\omega}}_T\|_2^2 T \right] + \mathcal{O}(T^2), \tag{S.3a}$$

where

$$\mathbf{G}_F[\hat{\dot{\boldsymbol{\omega}}}_T] := \frac{1}{2\|\nabla F[\mathbf{w}(0)]\|_2} \left\langle \hat{\dot{\boldsymbol{\omega}}}_T, \nabla^2 F[\mathbf{w}(0)] \hat{\dot{\boldsymbol{\omega}}}_T \right\rangle. \tag{S.3b}$$

### Decomposition of local task difficulty

In the main text $\mathbf{n}_2$ and $\mathbf{n}_3$ correspond to different sources of task-irrelevant synaptic changes (see equation 4 ). We treat them as random variables arising from some unknown probability distribution. Since they are generated by task-independent processes, they are uncorrelated with derivatives of the task error. Therefore

$$\mathbb{E}\left[ \left\langle \nabla F[\mathbf{w}(0)], \nabla^2 F[\mathbf{w}(0)] \mathbf{n}_i \right\rangle \right] = 0 \quad \text{for } i \in \{2, 3\}. \tag{S.4a}$$

We also assume that $\mathbb{E}[n_{2,i} n_{3,i}] = 0$, for any component $i$, i.e. the two sources of plasticity are uncorrelated with each other. This follows from the fact that $\mathbf{n}_3$ is the result of a white-noise process evolving at each synapse, and is thus uncorrelated with any random variable that is not itself derived from the same white-noise process. A consequence of this assumption is that

$$\mathbb{E}\left[ \left\langle \mathbf{n}_2, \nabla^2 F[\mathbf{w}(0)] \mathbf{n}_3 \right\rangle \right] = 0. \tag{S.4b}$$

We now substitute our expanded expression (see equation 5 of the main text)

$$\dot{\boldsymbol{\omega}}_T = -\gamma_1 \nabla \hat{F}[\mathbf{w}(0)] + \gamma_2 \hat{\mathbf{n}}_2 + \gamma_3 \sqrt{\frac{N}{T}} \hat{\mathbf{n}}_3. \tag{S.5}$$

for the interpolated synaptic velocity into our formula for the local task difficulty $\mathbf{G}_F[\hat{\dot{\boldsymbol{\omega}}}_T]$. We can use equations (S.4) to simplify the consequent expression, and thereby get

$$\mathbb{E}[\mathbf{G}_F[\hat{\dot{\boldsymbol{\omega}}}_T]] = \gamma_1^2 \mathbf{G}_F^1[\mathbf{w}(0)] + \gamma_2^2 \mathbf{G}_F^2[\hat{\dot{\boldsymbol{\omega}}}_T] + \gamma_3^2 \frac{N}{T} \mathbf{G}_F^3[\hat{\dot{\boldsymbol{\omega}}}_T], \tag{S.6}$$

where

$$\mathbf{G}_F^1[\mathbf{w}(0)] = \frac{1}{2\|\nabla F[\mathbf{w}(0)]\|_2}\Big\langle \nabla \hat{F}[\mathbf{w}(0)], \nabla^2 F[\mathbf{w}(0)]\nabla \hat{F}[\mathbf{w}(0)]\Big\rangle \tag{S.7a}$$

$$\mathbf{G}_F^2[\dot{\boldsymbol{\omega}}_T] = \frac{1}{2\|\nabla F[\mathbf{w}(0)]\|_2}\left\langle \hat{\mathbf{n}}_2, \nabla^2 F[\mathbf{w}(0)]\hat{\mathbf{n}}_2\right\rangle \tag{S.7b}$$

$$\mathbf{G}_F^3[\dot{\boldsymbol{\omega}}_T] = \frac{1}{2\|\nabla F[\mathbf{w}(0)]\|_2}\left\langle \hat{\mathbf{n}}_3, \nabla^2 F[\mathbf{w}(0)]\hat{\mathbf{n}}_3\right\rangle \tag{S.7c}$$

The independence of $\mathbf{n}_2$ and $\mathbf{n}_3$ from $\nabla^2 F[\mathbf{w}]$ allows us to further simplify the expressions for $\mathbf{G}_F^2$ and $\mathbf{G}_F^3$. Specifically, we can write $\mathbf{n}_i$ as

$$\mathbf{n}_i = \sum_{j=1}^{N} c_j v_j,$$

where $v_j$ denotes the $j^{th}$ eigenvector of $\nabla^2 F[\mathbf{w}]$. Independence of $\mathbf{n}_i$ from $\nabla^2 F[\mathbf{w}]$ implies that $\mathbb{E}[c_j] = \mathbb{E}[c_k]$ for any $j, k \in \{1, \dots, N\}$. This gives

$$\mathbb{E}\left\langle \hat{\mathbf{n}}_i, \nabla^2 F[\mathbf{w}(0)]\hat{\mathbf{n}}_i\right\rangle = \frac{Tr(\nabla^2 F[\mathbf{w}(0)])}{N},$$

the mean of the eigenvalues of $\nabla^2 F[\mathbf{w}(0)]$. With this, equation (S.6) becomes

$$\mathbb{E}\big[\mathbf{G}_F[\dot{\boldsymbol{\omega}}_T]\big] = \gamma_1^2 \mathbf{G}_F^1[\mathbf{w}(0)] + \frac{Tr(\nabla^2 F[\mathbf{w}(0)])}{2\|\nabla F[\mathbf{w}(0)]\|_2}\left[\frac{\gamma_2^2}{N} + \frac{\gamma_3^2}{T}\right]. \tag{S.8}$$

### 0.1 Learning from a distribution of inputs

We validated the analytic predictions of the main paper with numerical simulations. In order to do so, we constructed learning tasks for which the true gradient of task error was available (see Methods), and generated learning rules by corrupting this gradient with preset amounts of noise. In this section, we consider a common scenario that prevents exact calculation of the task gradient, but that corrupts any calculation with a degree of noise independent of network size. Specifically, tasks learn an input-output mapping over a continuous probability distribution of inputs, rather than a finite number of inputs (the case considered in numerical simulations accompanying the main text). Over each learning cycle, only a single input is drawn from the distribution. The gradient of task error can therefore only be calculated with respect to this single input, introducing an unavoidable degree of noise in any learning rule. This scenario is commonly called ‘stochastic gradient descent’.

The noise in a learning rule attributable to subsampling of inputs is independent of network size. It therefore enters the $\gamma_2$ term of weight change over each learning cycle. The amount by which it increases $\gamma_2$ is not easily calculable, and depends on both the task, and the distribution of inputs. We do not attempt this calculation. However, we do know that the value of $\gamma_2$ arising from subsampling will be shared between the small and large networks. Therefore, if we set $\gamma_3 = 0$, then increasing the size of the network will always improve learning rate / steady state performance. This is illustrated in Figure 1.

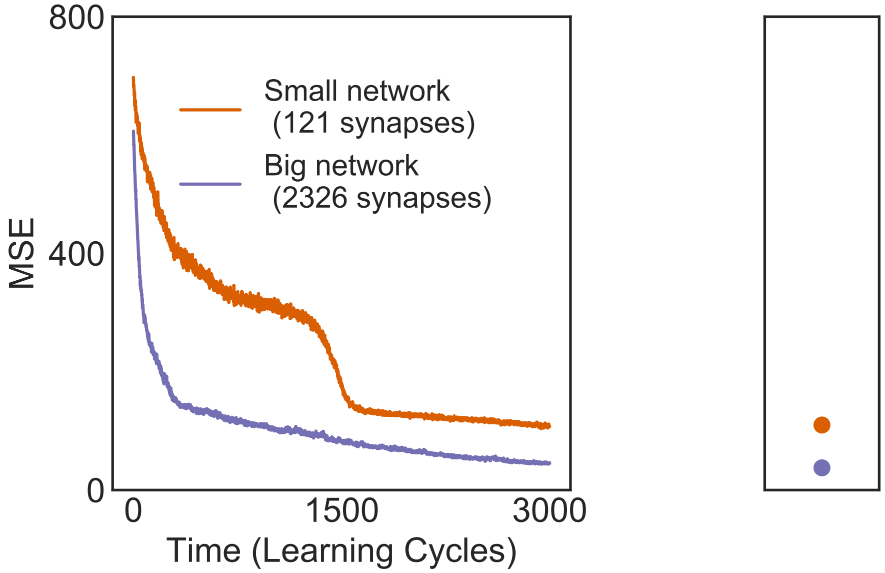


Figure 1: Dynamics of learning in a situation where the true task-error gradient cannot be computed, thus adding noise to the learning rule. Comparison of big and small networks. Both networks have the same number of inputs (10) and outputs (10), and learn the same fixed mapping from a teacher network. Over each learning cycle, a single input is drawn from a unit Gaussian distribution. The gradient of task error is calculated only with respect to this input (i.e. stochastic gradient descent). The task error is estimated by evaluating task error over 1000 Gaussian inputs. **Left**: Estimated task error is plotted over time for the big and small networks. **Right**: Mean steady state error after 3000 learning cycles, over 12 iterations. Error bars are invisible due to their small size: A single standard deviation from the mean is $[1.67858973, 1.0943906]$ for the small and big networks respectively.

### Task-relevant Plasticity

Recall that $\dot{\boldsymbol{\omega}}_T$ represents the vector of synaptic weight changes over a time interval $[0, T]$. A learning rule must determine both the speed and direction of synaptic plasticity, and thus $\dot{\boldsymbol{\omega}}_T$, over this time interval, using any task-related information is possesses. Suppose that the speed $\|\dot{\boldsymbol{\omega}}_T\|_2$ of this change was fixed at an arbitrary level. It remains for the learning rule to determine a direction

$\dot{\boldsymbol{\omega}}_T$ of synaptic plasticity over the time interval. If the objective of the learning rule was to minimise $F[\mathbf{w}(0) + \dot{\boldsymbol{\omega}}_T]$, the task error at time $t$, which direction should it pick?

First note that we can use a Taylor expansion to express the change in task error over $[0, T]$ as

$$F[\mathbf{w}(0) + T\dot{\boldsymbol{\omega}}_T] - F[\mathbf{w}(0)] = T\|\dot{\boldsymbol{\omega}}_T\|_2\langle\hat{\dot{\boldsymbol{\omega}}}_T, \nabla F[\mathbf{w}(0)]\rangle + T^2\|\dot{\boldsymbol{\omega}}_T\|_2^2\langle\hat{\dot{\boldsymbol{\omega}}}_T, \nabla^2 F[\mathbf{w}(0)]\hat{\dot{\boldsymbol{\omega}}}_T\rangle + \mathcal{O}[T^3\|\dot{\boldsymbol{\omega}}_T\|_2^3]. \tag{S.9}$$

Maximising the degree of learning involves minimising this quantity.

In the main text, we consider learning rules that have (at best) information on the gradient $\nabla F[\mathbf{w}(0)]$, but not on higher order derivatives $\nabla^{(n)}F[\mathbf{w}(0)]$. These are known as first-order learning rules. Given such information, a learning rule can only attempt to minimise the first term in (S.9). Hence, the optimal direction of plasticity would be the solution of

$$\begin{aligned}\hat{\dot{\boldsymbol{\omega}}}_T^{*,1} &= \min_{\hat{x}} T\|\dot{\boldsymbol{\omega}}_T\|_2\langle\hat{x}, \nabla F[\mathbf{w}(0)]\rangle : \qquad \|\hat{x}\|_2 = 1 \\ &= -\nabla\hat{F}[\mathbf{w}(0)].\end{aligned}$$

Indeed we define $-\hat{F}[\mathbf{w}(0)]$ (i.e. $\hat{\dot{\boldsymbol{\omega}}}_T^{*,1}$) as the direction of task-relevant plasticity in the main text. The expected correlation of additional, task-irrelevant plasticity with the Hessian $\nabla^2 F[\mathbf{w}(0)]$ is taken as zero, as must be true of any learning rule without access to information on the Hessian.

Learning rules that additionally have information on $\nabla^2 F[\mathbf{w}(0)]$ are known as second order learning rules. Given the additional information, $-\nabla\hat{F}[\mathbf{w}(0)]$ is no longer the best direction of plasticity. Instead, the optimal direction would minimise the first two terms of (S.9), and so would be

$$\hat{\dot{\boldsymbol{\omega}}}_T^{*,2} = \min_{\hat{x}} T\|\dot{\boldsymbol{\omega}}_T\|_2\langle\hat{x}, \nabla F[\mathbf{w}(0)]\rangle + T^2\|\dot{\boldsymbol{\omega}}_T\|_2^2\langle\hat{x}, \nabla^2 F[\mathbf{w}(0)]\hat{x}\rangle : \qquad \|\hat{x}\|_2 = 1.$$

Whereas $\hat{\dot{\boldsymbol{\omega}}}_T^{*,1}$ was a steepest descent direction of the task error, $\hat{\dot{\boldsymbol{\omega}}}_T^{*,2}$ looks for directions that balance immediate descent with both downward curvature of the task error, and the time $T$ until direction can be updated. Intuitively, a descent direction that is downwardly curved is likely to remain a descent direction as the weights change along it.

The solution to this minimisation has a complicated analytic form that is dependent upon both $T$ and $\|\dot{\boldsymbol{\omega}}_T\|_2$. In order to incorporate second order learning rules in the main text, we would have to change the direction of task-relevant plasticity from $\hat{\dot{\boldsymbol{\omega}}}_T^{*,1}$ to $\hat{\dot{\boldsymbol{\omega}}}_T^{*,2}$. How would using $\hat{\dot{\boldsymbol{\omega}}}_T^{*,2}$ as the direction of task-relevant plasticity change the results of the main text? All workings in the paper would have to replace the original formula (see equation (S.5)) for $\dot{\boldsymbol{\omega}}_T$ with

$$\dot{\boldsymbol{\omega}}_T = -\gamma_1\hat{\dot{\boldsymbol{\omega}}}_T^{*,2} + \gamma_2\hat{\mathbf{n}}_2 + \gamma_3\sqrt{\frac{N}{T}}.$$

This is tractable, but technically cumbersome. Of particular note is that

$$\langle \dot{\boldsymbol{\omega}}_T^{*,2}, \nabla^2 F[\mathbf{w}(0)] \dot{\boldsymbol{\omega}}_T^{*,2} \rangle \leq \langle \dot{\boldsymbol{\omega}}_T^{*,1}, \nabla^2 F[\mathbf{w}(0)] \dot{\boldsymbol{\omega}}_T^{*,1} \rangle.$$

Consequently, the formula for (S.7a), the component of local task difficulty attributable to task-relevant plasticity, would decrease, as we would get

$$\mathbf{G}_F^1[\mathbf{w}(0)] = \frac{1}{2\|\nabla F[\mathbf{w}(0)]\|_2} \left\langle \dot{\boldsymbol{\omega}}_T^{*,2}, \nabla^2 F[\mathbf{w}(0)] \dot{\boldsymbol{\omega}}_T^{*,2} \right\rangle.$$

Qualitatively, the results of the paper would not change in that the relationship between task-irrelevant plasticity and network size would be preserved. Task-irrelevant plasticity would still be uncorrelated with the derivatives of task error. So we would arrive at an identical equation (S.8), except for the changed formula of $\mathbf{G}_F^1[\mathbf{w}(0)]$.

Note that we could analogously consider $n^{th}$ order learning rules, for arbitrary $n$, by taking the direction of task-relevant plasticity as $\dot{\boldsymbol{\omega}}_T^{*,n}$, where this direction minimises the first $n$ terms of the Taylor expansion (S.9). A learning rule with access to derivatives of all orders $n$, given an analytic task-error function, is equivalent to a learning rule that knows *a priori* the optimal weight configuration of the network, and induces synaptic plasticity in a direction pointing directly towards this optimal configuration. We provide this observation for intuition only, and do not suggest that such a learning rule would realistically exist in a biological context.

### Learning in a Linear Network

We consider the simple case of a linear network undertaking supervised learning with a quadratic error function. The general (nonlinear) case is considered subsequently. Inputs $u$ are drawn from some space $\mathcal{U} \subseteq \mathbb{R}^i$ with probability $\mathbb{P}(u)$. For each input $u$, there is an optimal output (i.e. label) $y^*(u) \in \mathbb{R}^o$.

The network linearly transforms any input into an output $y = Wu$, for a matrix $W \in \mathbb{R}^{oi}$ of synaptic weights. Note the cosmetic difference between the matrix $W$, and previous representations of the synaptic weights, which took the form of a vector $\mathbf{w} \in \mathbb{R}^N$. These representations are interchangeable, as we can reshape $W$ into a vector $\mathbf{w}$, and take $N = oi$. Indeed, we will switch between the matrix and vector representations of the weights in the ensuing discussion. The input-dependent error of the network, for any particular input $u \in \mathcal{U}$, is taken as

$$F(W, u) = \|y^*(u) - Wu\|_2^2.$$

Meanwhile, the overall network error integrates input-dependent error over the probability distribution $\mathbb{P}(u)$ of inputs, i.e.

$$F[W] = \int_{u \in \mathcal{U}} F[W, u] \;\; \mathbb{P}(u) \; du.$$

Even if $y^*(u)$ were known for each $u \in \mathcal{U}$, calculation of $F[W]$ would require passage of every input $u \in \mathcal{U}$ through the network. In a biologically realistic setting, $y^*(u)$ would not be known, and both $F[W]$ and its gradient would have to be estimated, using a subset of inputs. So no learning rule minimising the error $F[W]$ could have uncorrupted access to $\nabla F[W]$.

Now let us lift the learning problem into a higher dimensional setting. We will consider a matrix $W'$ with $c_1 i$ inputs and $c_2 o$ outputs, for some constants $c_1$, $c_2 > 1$. We will define the total number of weights as $\tilde{N} = c_1 i c_2 o$. We take the transformation $u' = Bu \in \mathbb{R}^{c_1 i}$, where $B \in \mathbb{R}^{c_1 i \times i}$ is an arbitrary semi-orthogonal matrix. (i.e. it satisfies $B^T B = \mathbb{I}_i$). Geometrically, $B$ therefore represents the composition of a projection into the higher dimensional space $\mathbb{R}^{c_1 i}$ with a rotation. Note that this is an invertible mapping: if $u' = Bu$ then $B^T u' = u$. Similarly, we can take $y'^*(u') = Dy^*(u) \in \mathbb{R}^{c_2 o}$, where $D^T D = \mathbb{I}_o$.

The expanded neural network with weights $W' \in \mathbb{R}^{c_1 o \times c_2 i}$ has to learn the same mapping as the original, but with respect to the higher dimensional inputs. So the network receives inputs $u' \in B\mathcal{U}$, and transforms them to outputs $y' = W'u'$, with input-dependent error

$$F'[W', u'] = \|y'^*(u') - W'u'\|_2^2. \tag{S.10a}$$
$$= \|Dy^*(u) - W'Bu\|_2^2 \text{ for some } u : u' = Bu \tag{S.10b}$$
$$= \|y^*(u) - D^T W'Bu\|_2^2 \text{ by semi-orthogonality of } D. \tag{S.10c}$$

Let us randomly draw a state $W' \in \mathbb{R}^{c_1 o \times c_2 i}$ of the bigger network. For instance, each element of $W'$ can be drawn from a Gaussian distribution. Regardless of $W'$, equations (S.10) show us that $W = D^T W' B \in \mathbb{R}^{oi}$ defines a state of the original network satisfying

$$F'[W'] = F[W].$$

If we consider the synaptic weight matrices $W'$ and $W$ as vectors $\mathbf{w}' \in \mathbb{R}^{\tilde{N}}$ and $\mathbf{w} \in \mathbb{R}^N$, then the mapping can be represented as

$$\mathbf{w} = H\mathbf{w}', \quad HH^T = \mathbb{I}_N.$$

By the chain rule:

$$H\nabla F'[\mathbf{w}'] = \nabla F[\mathbf{w}],$$
$$H\nabla^2 F'[\mathbf{w}']H^T = \nabla^2 F[\mathbf{w}].$$

Semi-orthogonality of $H$ implies that it has $N$ singular values with value one, and $\tilde{N} - N$ singular values with value zero. We take the heuristic that $\mathbf{w}'$ should project approximately equally onto each of the associated singular vectors. This is reasonable given that we took a random choice of $W'$, and implies

$$\frac{\|\nabla F'[\mathbf{w}']\|_2}{\|\nabla F[\mathbf{w}]\|_2} \approx \frac{\tilde{N}}{N} = c_1 c_2 \tag{S.11}$$

Meanwhile the quadratic nature of the error function implies $\nabla^2 F'[\mathbf{w}]$ is constant. The trace of this matrix can be calculated explicitly as

$$Tr(\nabla^2 F'[\mathbf{w}']) = (c_2 o)^2 \int_{u \in \mathcal{U}} \|u\|_2^4 \ \ \mathbb{P}(u). \tag{S.12}$$

We will also assume that $\nabla \hat{F'}[\mathbf{w}]$ (which is a normalised vector) projects approximately equally onto the different eigenvectors of $\nabla^2 F'[\mathbf{w}']$. The latter is fixed, regardless of $\mathbf{w}'$, whereas the former is a linear function of the (randomly chosen) $\mathbf{w}$, which justifies the approximation. In this case, (S.12) implies

$$\nabla \hat{F'}[\mathbf{w}']^T \ \nabla^2 F'[\mathbf{w}'] \ \nabla \hat{F'}[\mathbf{w}'] \approx (c_2)^2 \nabla \hat{F}[\mathbf{w}]^T \ \nabla^2 F[\mathbf{w}] \ \nabla \hat{F}[\mathbf{w}]. \tag{S.13}$$

Bringing together equations (S.11), (S.13), and the formula (S.7a) for $\mathbf{G}_F^1$, we see that

$$\mathbf{G'}_{F'}^1[\mathbf{w}'] \approx \frac{(c_2)^2}{c_1 c_2} \mathbf{G}_F^1[\mathbf{w}] = \frac{c_2}{c_1} \mathbf{G}_F^1[\mathbf{w}]. \tag{S.14a}$$

Similarly

$$\frac{Tr(\nabla^2 F'[\mathbf{w}'])}{\|\nabla F'[\mathbf{w}']\|_2} \approx \frac{c_2}{c_1} \frac{Tr(\nabla^2 F[\mathbf{w}])}{\|\nabla F[\mathbf{w}]\|_2}. \tag{S.14b}$$

Now recall the learning rate equation, which can be rewritten as

$$k = \frac{-\|\nabla F[\mathbf{w}(0)]\|_2}{F[\mathbf{w}(0)]} \Big[ -\gamma_1 + \boldsymbol{\delta} \Big] + \mathcal{O}(T^2).$$
$$\boldsymbol{\delta} = \mathbf{G}_F[\dot{\hat{\boldsymbol{\omega}}}_T] \|\dot{\boldsymbol{\omega}}_T\|_2^2 T,$$

with $\mathcal{O}(T^2) \equiv 0$ for a quadratic error function such as we have. As long as we preserve the ratio $\frac{c_2}{c_1}$, equations (S.14) allow us to consider $N$ as an independent parameter in the equation for $\boldsymbol{\delta}$. This allows us to optimise steady state error of the network by changing $N$. To see how, suppose the network has reached steady state error, i.e $\mathbb{E}[k] = 0$. If we decreased $\boldsymbol{\delta}$, then $\mathbb{E}[k]$ would increase, and the network would learn further. Therefore, to derive the optimal $N^*$, we should minimise the expression for $\boldsymbol{\delta}$ in $N$. We differentiate $\boldsymbol{\delta}$ in $N$, and note that stationary points satisfy the equation:

$$N^2 \left[ \gamma_1^2 C T + \gamma_3^2 \right] = \frac{T^2 \gamma_2^2}{\gamma_3^2} \left( \gamma_1^2 + \gamma_2^2 \right) \tag{S.15}$$

$$\text{where } C = \frac{\Big\langle \nabla \hat{F}[\mathbf{w}(0)], \nabla^2 F[\mathbf{w}(0)] \nabla \hat{F}[\mathbf{w}(0)] \Big\rangle}{Tr(\nabla^2 F[\mathbf{w}(0)])}. \tag{S.16}$$

For $\gamma_2 \neq 0$, this implies the existence of two stationary points differing only in sign. Since $\lim_{N \to \infty} \boldsymbol{\delta} = \infty$, and $\lim_{N \to -\infty} \boldsymbol{\delta} = -\infty$, the positive stationary point is necessarily a global minimum of $\boldsymbol{\delta}$. So this stationary point defines $N^*$. We have

$$N^* = \frac{T\gamma_2}{\gamma_3}\left[\sqrt{\frac{1+\frac{\gamma_2^2}{\gamma_1^2}}{CT+\frac{\gamma_3^2}{\gamma_1^2}}}\right]. \tag{S.17}$$

Note that $C$ is unknown, and depends on the particular weight configuration of the network. However, we can take the heuristic $C \approx \frac{1}{N^*}$. This heuristic is exact if the gradient $\nabla \hat{F}[\mathbf{w}(0)]$ projects equally onto each of the eigenvalues of the $\nabla^2 \hat{F}[\mathbf{w}(0)]$. This in turn would make the numerator of $C$ equal the mean eigenvalue, i.e. $\frac{Tr(\nabla^2 \hat{F}[\mathbf{w}(0)])}{N}$. This heuristic should hold on average over time, given that the Hessian $\nabla^2 \hat{F}[\mathbf{w}(0)]$ is fixed throughout learning due to the quadratic error function, while, the direction $\nabla^2 \hat{F}[\mathbf{w}(0)]$ depends completely on $\mathbf{w}(0)$, which changes over each learning cycle in a direction independent of $\nabla^2 \hat{F}[\mathbf{w}(0)]$.

We now have an expression for the optimal network size. Critical to our derivation was the ability to take an arbitrary state $W'$ of the larger network, and find a corresponding state $W$ of the nominal network that was linked to $W'$ via a transformation that preserved task error. This will not be the case in subsequent, more general examples.

In summary, we have considered linear networks with quadratic error functions learning a random mapping. We have provided a transformation that increases the size of the linear network, and embeds the random mapping in a higher dimensional space. The existence of this transformation assures us that the higher-dimensional learning problem should have a lower task difficulty. Given a synaptic learning rule with known levels of gradient information, systematic error, and intrinsic noise, we can predict a priori what the optimal dimensionality of the linear network should be, if we want the network to minimise steady state error.

### Learning in a nonlinear, feedforward network

We now extend to the case of multilayer, feedforward networks with nonlinearities, before finally considering the general case. We will consider a static nonlinearity $\sigma : \mathbb{R} \to \mathbb{R}$, which can be overloaded to give a function $\underline{\sigma} : \mathbb{R}^m \to \mathbb{R}^m$ representing the elementwise application of $\sigma$ to a vector of length $\mathbb{R}^m$, for arbitrary $m \in \mathbb{N}$. The $k^{th}$ layer of the network transforms any input $u^{(k-1)}$ from the previous layer into an output of the form

$$y^{(k)} = \underline{\sigma}(W^{(k)}u^{(k-1)}).$$

Again, input-dependent error is taken as the squared deviation between the network output and some optimal output $y^*(u)$. We take $W$ as the concatenation of the weight matrices $W^{(k)}$ at each layer $k$. For instance, in the case of a two-layer network, we have

$$F(W, u) = \|y^*(u) - \sigma\big(W^{(2)}\sigma(W^{(1)}u)\big)\|_2^2.$$

In the linear example, we considered a random state (i.e. weight configuration) of the expanded network, and transported it to an equivalent state of the nominal network via an error-preserving transformation. In the present example this is not possible, as the expanded network can exhibit a fundamentally higher-dimensional set of behaviours. Instead, we will construct an error-preserving transformation $\phi$ in the opposite direction, from states of the nominal network to states of the larger network. This transformation will essentially add neurons to the network so that gradient information is distributed over a larger number of neurons. Our analysis will quantify the effect of this network expansion on local task difficulty, and hence determine an optimal network size. Technically, it will only hold for network states of the expanded network that reside in the image of the mapping $\phi$. However, we subsequently justify why extrapolating from these network states to generic states is reasonable.

For the synaptic weight matrix $W^{(k)} \in \mathbb{R}^{o_k i_k}$ of the $k^{th}$ layer, let us choose a transformation $\phi^k(W^{(k)}) = D^{(k)}W^{(k)}(B^{(k)})^T \in \mathbb{R}^{o'_k \times i'_k}$ (from the original matrix to a bigger matrix with dimensionality $o'_k \times i'_k$) that preserves task error. We will take

$$B^{(k)} = [\mathbb{I}_{i_k}, 0_{i'_k - i_k}]$$
$$D^{(k)} = [\mathbb{I}_{o_k}, 0_{o'_k - o_k}].$$

For the input layer, we demand that $i'_1 = i_1$ so that the number of inputs is preserved and $B^{(1)}$ is the identity matrix. For the output ($d^{th}$) layer, we will similarly demand that $o'_d = o_d$ so that the output dimensionality of the network is preserved. Intermediate layers in the transformed network have, by the above construction, greater (or equal) numbers of neurons as compared to the nominal network. Regardless of the nonlinearity $\sigma$ (which may not satisfy $\sigma(0) = 0$, and indeed does not for sigmoidal nonlinearities as used in numerical examples of the paper), this transformation satisfies

$$F'[\phi(W)] = F[W].$$

Here we have essentially added synapses to the network with weight zero, in such a way that they contribute nothing to the input-output properties of the network. However as soon as their weights deviate from zero, this is no longer the case. So they do contribute to the gradient $\nabla F'[\phi(W)]$. If we represent the concatenated weight matrices of the network as a long vector $\mathbf{w}$, the described transformation can be effected by a function $\phi(\mathbf{w}) = A\mathbf{w} \in \mathbb{R}^{\tilde{N}}$, where $\tilde{N}$ is the number of weights in the expanded network, and where the matrix $A$ satisfies

$A^T A = \mathbb{I}_N$. Using the chain rule, we have, for any $\mathbf{w}^* \in \mathbb{R}^N$:

$$A^T \nabla F'[\phi(\mathbf{w}^*)] = \nabla F[\mathbf{w}^*] \tag{S.18a}$$

$$A^T \nabla^2 F'[\phi(\mathbf{w}^*)] A = \nabla^2 F[\mathbf{w}^*]. \tag{S.18b}$$

We will use equations (S.18) to somewhat constrain the learning rate. First note that

$$\|\nabla F'[\phi(\mathbf{w}^*)]\|_2 \geq \|\nabla F[\mathbf{w}^*]\|_2 \tag{S.19a}$$

through equation (S.18a) and semi-orthogonality of $A$. Heuristically, we can say that

$$\frac{\|\nabla F'[\phi(\mathbf{w}^*)]\|_2}{\|\nabla F[\mathbf{w}^*]\|_2} \approx \frac{\tilde{N}}{N}. \tag{S.19b}$$

This would be true if $\|\nabla F'[\phi(\mathbf{w}^*)]\|_2$ projected onto each of the eigenvectors of $AA^T$ equally, given that $AA^T$ has $N$ eigenvalues of magnitude one, and $\tilde{N} - N$ of magnitude zero. In the same spirit, we take the heuristic

$$Tr(A^T \nabla^2 F'[\phi(\mathbf{w}^*)] A) \equiv Tr(\nabla^2 F'[\phi(\mathbf{w}^*)] AA^T) \approx \frac{\tilde{N}}{N} Tr(\nabla^2 F[\mathbf{w}^*]). \tag{S.19c}$$

The approximation similarly holds if the sum of the columns of $\nabla^2 F'[\phi(\mathbf{w}^*)]$ projects equally onto each of the eigenvectors of $AA^T$. Note that this approximation would not be reasonable in the previous linear example with a quadratic error function. In that case $\nabla^2 F'$ was constant over learning. Since $A$ is also constant, if one encountered a situation where e.g. $\nabla^2 F'$ projected much more strongly onto eigenvectors of $AA^T$ with nonzero eigenvalues, then that situation would hold over the entire course of learning, since it related two constant matrices. In the current, nonlinear example, by contrast, the error function is non-quadratic, and thus $\nabla^2 F'$ changes over time. It has columns that continually shift direction over the course of learning. On average, it is therefore reasonable to assume that their sum projects onto $AA^T$ in the manner hypothesised.

Finally, expansion of equations (S.18) gives

$$\begin{aligned} \left\langle \nabla F[\mathbf{w}^*], \nabla^2 F[\mathbf{w}^*] \nabla F[\mathbf{w}^*] \right\rangle &= \left\langle A^T \nabla F'[\phi(\mathbf{w}^*)], A^T \nabla^2 F'[\phi(\mathbf{w}^*)] AA^T \nabla F'[\phi(\mathbf{w}^*)] \right\rangle. \\ &\approx \left(\frac{N}{\tilde{N}}\right)^2 \left\langle \nabla F'[\phi(\mathbf{w}^*)], \nabla^2 F'[\phi(\mathbf{w}^*)] \nabla F'[\phi(\mathbf{w}^*)] \right\rangle, \end{aligned}$$

where we again assume uniformity of projection of $\nabla F'$ onto the eigenalues of $AA^T$ to make the approximation. Normalising the LHS and RHS of the above equation by $\|\nabla F[\mathbf{w}^*]\|_2^2$ and $\|\nabla F'[\mathbf{w}^*]\|_2^2$ respectively, equation (S.19b) implies

$$\left\langle \nabla \hat{F}[\mathbf{w}^*], \nabla^2 F[\mathbf{w}^*] \nabla \hat{F}[\mathbf{w}^*] \right\rangle \approx \left\langle \nabla \hat{F}'[\phi(\mathbf{w}^*)], \nabla^2 F'[\phi(\mathbf{w}^*)] \nabla \hat{F}'[\phi(\mathbf{w}^*)] \right\rangle. \tag{S.19d}$$

This, along with (S.19c), allows us to estimate how $\mathbf{G}_F^1$ grows with network size. We can then use equations (S.19) to estimate how local task difficulty varies with the number of synapses. Given $N$ synapses in the original network, and $\tilde{N}$ in the expanded network, we get

$$\mathbb{E}[\mathbf{G'}_{F'}[\phi(\dot{\boldsymbol{\omega}}_T)]] \approx \frac{N}{\tilde{N}}\gamma_1^2\mathbf{G}_F^1[\mathbf{w}(0)] + \frac{Tr(\nabla^2 F[\mathbf{w}(0)])}{2\|\nabla F[\mathbf{w}(0)]\|_2}\left[\frac{\gamma_2^2}{\tilde{N}} + \frac{\gamma_3^2}{T}\right],$$

where $\mathbf{G'}_{F'}$ is the local task difficulty associated with the direction $\phi(\dot{\boldsymbol{\omega}}_T)$ in the expanded network, given an initial synaptic configuration of $\phi[\mathbf{w}(0)]$. As we did in the linear case (see equation (S.17)), we can differentiate the expression for $\boldsymbol{\delta}$ in $N$, to find an $N^*$ that optimises learning rate. We get

$$N^* = \frac{T\gamma_2}{\gamma_3^2}\left[\sqrt{\Big(1 + \frac{\gamma_1^2 CN}{\gamma_2^2}\Big)\left(\gamma_1^2 + \gamma_2^2\right)}\right], \tag{S.20a}$$

$$\approx \frac{T\gamma_2}{\gamma_3^2}\left[\sqrt{\Big(1 + \frac{\gamma_1^2 N}{\gamma_2^2 N^*}\Big)\left(\gamma_1^2 + \gamma_2^2\right)}\right] \tag{S.20b}$$

where $N$ is the number of synapses in the original network, and $C$ is as defined in equation (S.16). Also as in equation (S.16), we take the approximation $C = \frac{1}{N^*}$. This leads to (S.20a) which can be expressed as a cubic equation in $N^*$ with a single positive root, and thus solved.

We have now considered a multilayer feedforward network with static nonlinearities at each neuron. We have constructively showed how any state of this layer can be embedded in a higher-dimensional layer in a manner that quantifiably lowers task difficulty. One issue with the analysis is that we only consider the task difficulty of weight vectors in the image of $\phi$ (for the expanded network). This is a low-dimensional subset of the overall set of attainable weight vectors. For the analysis to be valid, we must assume that the expected local task difficulty is not much lower for weights in the image of $\phi$, than for other weights with the same task error. If this were not true, then the local learning rate of the expanded network would jump whenever the weight vector of the network was in the image of $\phi$. Given that $\phi$ is constructed with respect to the network architecture, but not the specific task being learnt, we find this unlikely in general.

The mathematical analysis we conducted can be used as a template to conduct size-increasing transformations on more general neural network architectures. Suppose we have a generic neural network with synaptic weights $\mathbf{w}$. Suppose we add neurons to this generic neural network, and can fix the synaptic weights of the added neurons in such a way that the input-output properties of the network do not change. So the expanded network has weights $[\mathbf{w}, \tilde{\mathbf{w}}]$. Then the size-expanding neural network transformation can be expressed as a function $\phi$, with $\phi(\mathbf{w}) = [\mathbf{w}, \tilde{\mathbf{w}}]$. Since the input-output properties of the network do not change, we get

$$F'[\phi(\mathbf{w})] = F[\mathbf{w}].$$

Since $\phi(\mathbf{w}) = [\mathbf{w}, \tilde{\mathbf{w}}]$, we get

$$\big(\nabla\phi(\mathbf{w})\big)^T \big(\nabla\phi(\mathbf{w})\big) = \mathbb{I}_N.$$

At any particular weight configuration $\mathbf{w}^*$, we can locally define $\nabla\phi(w) = A\mathbf{w} + B$, for some $B$. Note that the above equation implies semi-orthogonality of $A$, which allows us to repeat the original analysis of the current section.
